# Placental extracellular vesicles from early-onset but not late-onset preeclampsia induce a pro-vasoconstrictive and anti-vasodilatory state in resistance arteries

**DOI:** 10.1101/2024.06.04.597453

**Authors:** Sien Yee Lau, Katie Groom, Colin L. Hisey, Qi Chen, Carolyn Barrett, Larry Chamley

## Abstract

**Background:** The human placenta releases large numbers of extracellular vesicles (EVs) into the maternal circulation throughout pregnancy. In preeclampsia, a hypertensive disorder of pregnancy, the number of EVs increases and the cargo they carry is altered. We investigated whether human placental EVs from pregnancies complicated by preeclampsia directly alter maternal vascular function, a hallmark of the disorder, and if EVs from early-onset or late-onset variants of preeclampsia have different effects on the vasculature.

**Methods:** Macro-EVs, micro-EVs and nano-EVs were isolated from cultured explants of human placentae from women with early-onset or late-onset preeclampsia, or from normotensive women. EVs were injected intravenously into pregnant mice and either at 30 minutes or 24 hours after injection, the mice were euthanized and the function of second order mesenteric arteries were assessed using wire myography.

**Results:** Placental EVs from pregnancies with early-onset preeclampsia enhanced vasoconstriction to PE, AngII, and ET-1 whilst impairing vasodilation to ACh and SNP in a time-dependent fashion, most prominently at 24 hours. In contrast, placental EVs from pregnancies with late-onset preeclampsia induced few differences compared to arteries taken from control mice injected with EVs from women with normotensive pregnancies.

**Conclusions:** To the best of our knowledge, this is the first comparison of vascular function after exposure to the full range of EVs produced by placentae from women with early-onset and late-onset preeclampsia and normotensive women. Placental EVs from early-onset preeclampsia demonstrated the ability to contribute to the development of the high-resistance haemodynamic profiles of women affected by early-onset preeclampsia.

## Introduction

Throughout pregnancy, the human placenta releases subcellular fragments called extracellular vesicles (EV) directly into the maternal circulation. EVs contain complex protein, lipid and nucleic acid cargos encased within phospholipid membranes. Most eukaryotic cells extrude two types of EV categorized loosely based on size, including micro-EVs (100-1000nm also called large EVs) and nano-EVs (<150nm, also called small EVs) (1). In addition to these two common types of EVs, the human placenta also uniquely extrudes macro-EVs (multinucleated structures up to 400µm in diameter). Macro-EVs are subcellular fragments of the unique multinucleated syncytiotrophoblast that lines the human placenta.

There are numerous reports demonstrating that EVs of various origins, such as plasma (2, 3), endothelial cells (4, 5), erythrocytes (6), lymphocytes (7, 8) and hepatocytes (9) can affect the response of resistance arteries to various vasodilators and vasoconstrictors. In pregnancy, the release of placental EVs directly into the maternal circulation adds an additional source of EVs which may have the potential to alter maternal vasculature. We have previously reported that EVs harvested from first trimester placental tissue explants can enhance vascular reactivity to the vasodilator acetylcholine 30 minutes after administration to mice *in vivo*, and that there is a shift in this response in the opposite direction at 24 hours (10). Others have also demonstrated the ability of placental EVs from term pregnancies to alter bradykinin-mediated vasodilation at 24 hours (11). This suggests placental EVs are able to directly modify the vasoactive response of maternal resistance arteries.

In healthy pregnancies, placental EVs are thought to be involved in the normal physiological adaptations to pregnancy (1). These effects are thought to be executed by various bioactive proteins carried by placental EVs, such as flt-1 (12, 13), eNOS (14) and neprilysin (15). Furthermore, placental EVs have also been shown to contain mRNA (16) and miRNA (17) which can be transferred into endothelial cells, giving the placental EVs the potential to exert longer term consequences on the target cells.

Preeclampsia is a hypertensive disorder of pregnancy with multi-organ involvement and a major cause of morbidity for mothers and babies (18). Preeclampsia is further sub-categorized into early-onset (developing <34 weeks of pregnancy) and late-onset (≥34 weeks), with the former being typically associated with poorer health outcomes (19). One hallmark of preeclampsia is systemic endothelial dysfunction, which is detectable weeks prior to the onset of clinical signs (20). While women with either early-onset or late-onset preeclampsia present clinically with higher systemic vascular resistance compared to normotensive women, women with early-onset, but typically not late-onset, preeclampsia also demonstrate lower cardiac output, suggesting a different hemodynamic profile between the two sub-types of preeclampsia (reviewed in 21).

There is increased release of placental EVs into the maternal circulation in preeclampsia (22–24) and the protein content of EVs from placentae of women with preeclampsia have also been reported to differ from that of placental EVs from normal pregnancies (25, 26). Given the ability of placenta-derived EVs to directly modulate the response of resistance arteries to vasoactive substances, we hypothesized that the altered nature of placental EVs in preeclampsia may induce a pro-constrictive response in arterial resistance vessels.

The effect on small artery function exhibited by EVs isolated directly from the human placenta in early-onset and late-onset preeclampsia has not been investigated. In this study, we used an established tissue explant model to isolate placental EVs to assess whether, individually, the three types of placental EVs from pregnancies with either early-or late-onset preeclampsia affect the vascular reactivity of resistance arteries differently from placental EVs from women with normotensive pregnancies. Specifically, we assessed the reactivity to angiotensin II (AngII), endothelin-1 (ET-1), acetylcholine (ACh) and sodium nitroprusside (SNP) due to the of the implication of the renin-angiotensin system (27), endothelin signaling (28, 29) and nitric oxide system (30, 31) in the development of the hypertension of preeclampsia. We also investigated whether these effects were different at 30 minutes, and 24 hours after circulating *in vivo*.

## Methods

### Collection of placentae and isolation of EVs

The collection and use of human placentae for this work was approved by a New Zealand Health and Disabilities Ethics Committee (approval number NXT1206057AM09). Placental EVs were isolated from placentae from pregnancies complicated by third trimester preeclampsia and normotensive pregnancies using a well-established tissue culture system (32) Preeclampsia was defined by the SOMANZ guidelines where early-onset preeclampsia is the development of clinical signs <34 weeks and late-onset preeclampsia ≥34 weeks of gestation (19). Women with known fetal chromosomal abnormalities, evidence of diabetes or autoimmune/inflammatory diseases were excluded from this study.

All women included in the early-onset and late-onset preeclampsia groups experienced proteinuria in addition to new-onset hypertension, with the early-preeclampsia group experiencing more diagnostic features (according to the SOMANZ guidelines (19)). Maternal characteristics and pregnancy outcomes are shown in Supplementary Table 1.

The maternal facing surface was dissected and discarded. Villous tissue explants of approximately 400 mg were dissected avoiding the basal plate and areas of excessive connective tissue. Excess blood was washed off the explants with sterile PBS and cultured overnight (16-24 hours) in NetWell™ inserts in 12 well plates in Advanced DMEM/F12 media containing 2% fetal bovine serum and 1% penicillin/streptomycin. The conditioned media was aspirated and centrifuged for 5 mins at 2000 x *g* to harvest macro-EVs. The supernatant was further centrifuged at 20,000 x g for 60 minutes to harvest the pelleted micro-EVs, and the supernatant centrifuged for 60 mins at 100,000 x g to collect pelleted nano-EVs. Contaminating red blood cells were removed from the macro-EV fraction by hypotonic lysis and CD45+ cells were removed by magnetic anti-CD45 Dynabeads (32). EVs were resuspended in PBS and stored at 4°C for up to four weeks until use.

The morphology and size of placental EVs micro– and nano-were confirmed by nanoparticle tracking analysis (NTA) and transmission electron microscopy (TEM) (details in the supplementary methods). Western blots for CD63+, calnexin-, cytokeratin 7+ and vimentin-nature of the placental EVs collected using our explant culture method has been published (33). The protein content of the EV preparations was quantified by bicinchoninic acid protein assay (Pierce™ BCA Protein Assay Kit, Thermofisher).

### Animal experiments

Animal experiments were approved by the University of Auckland Animal Ethics Committee (R002112). Pregnant CD-1 mice 8-11 weeks of age were administered a single intravenous injection of placental EVs via the tail vein on day 12.5 of gestation. Each intravenous injection comprised 100 µg protein of either micro-EVs or nano-EVs, or the macro-EVs isolated from 1.6 g of placental villous tissue suspended in 100 µL of sterile PBS. Mice were exposed to the EVs for either 30 minutes or 24 hours before they were euthanized by cervical dislocation. A summary of the experimental design and animal groups can be found in supplementary figure 1.

### Assessment of vascular reactivity by wire myography

Immediately following euthanasia, the intestinal and mesenteric tissues were dissected from the mice and placed in ice-cold physiological saline solution (PSS) preequilibrated with carbogen. Second order mesenteric arteries were dissected and mounted onto wire myograph chambers (DMT, Denmark) using 25 µm gold-coated tungsten wire. Throughout the experiment, the PSS in the chamber bath was heated to 37°C, bubbled with carbogen and replaced with fresh PSS every 20-30 minutes.

Prior to assessment of responses to vasoactive substances, the arteries were normalized to 13 kPa and a wake-up protocol was performed by challenging with 0.1 M K+ PSS, 10^-5^ M phenylephrine (PE), and 10^-5^ M acetylcholine (ACh) on 50% pre-constricted arteries. Arteries were only used if they responded to >90% relaxation when challenged with 10^-5^ M ACh confirming integrity of the endothelial layer of the artery. Vasoactive agents PE, ACh, sodium nitroprusside (SNP), angiotensin II (Ang II) and endothelin-1 (ET-1) (Sigma Aldrich, NZ) were added to reach cumulative concentrations ranging from 10^-11^ M to 10^-5^ M. Two arteries were mounted per mouse. Force exerted during contraction (and relaxation) were acquired using the Powerlab acquisition device and LabChart Acquisition Software (ADInstruments, NZ).

### Statistical analysis

To compare the responses to vasoactive substances between different arteries, the forces detected by the wire myograph were normalized. Contractile responses elicited by vasoconstrictors were normalized against the maximum stabilized contraction elicited by KPSS in the same vessel. Relaxation responses were normalized against the stable pre-constriction contractile response immediately prior to adding the vasodilator. Normalized contractile and relaxation responses were fitted with a 3-parameter non-linear regression using Prism (GraphPad Prism 8.0), with the least square regression and testing for significant differences using the extra sum of squares F test. A p-value <0.05 was deemed statistically significant.

## Results

### Characteristics of EVs collected from early-onset preeclampsia, late-onset preeclampsia and normotensive placentae

There were no obvious differences in the morphology of either the micro-EVs and the nano-EVs between early-onset preeclampsia, late-onset preeclampsia or normotensive placentae when examined by TEM (Supplementary Figure 2A). Likewise, the size distributions and modal sizes of the EVs were similar between the three conditions when assessed by NTA (Supplementary Figure 2B).

### Vascular responsiveness of resistance arteries 24 hours after injection of preeclamptic or normal placental EVs

#### Response to placental EVs from pregnancies with early-onset preeclampsia 24 hours after injection

All three subtypes of placental EVs isolated from pregnancies with early-onset preeclampsia increased the vascular reactivity of mesenteric arteries from pregnant mice injected intravenously to all three vasoconstrictors tested; PE, AngII and ET-1 (all p<0.05, blue circles, Figure 1), compared to mice injected with placental EVs from normotensive pregnancies (black squares, Figure 1). We were unable to fit the contraction responses to AngII in vessels from mice exposed to microvesicles using the 3-parameter non-linear regression and we were unable to determine whether the responses were different between groups.

**Figure 1.**
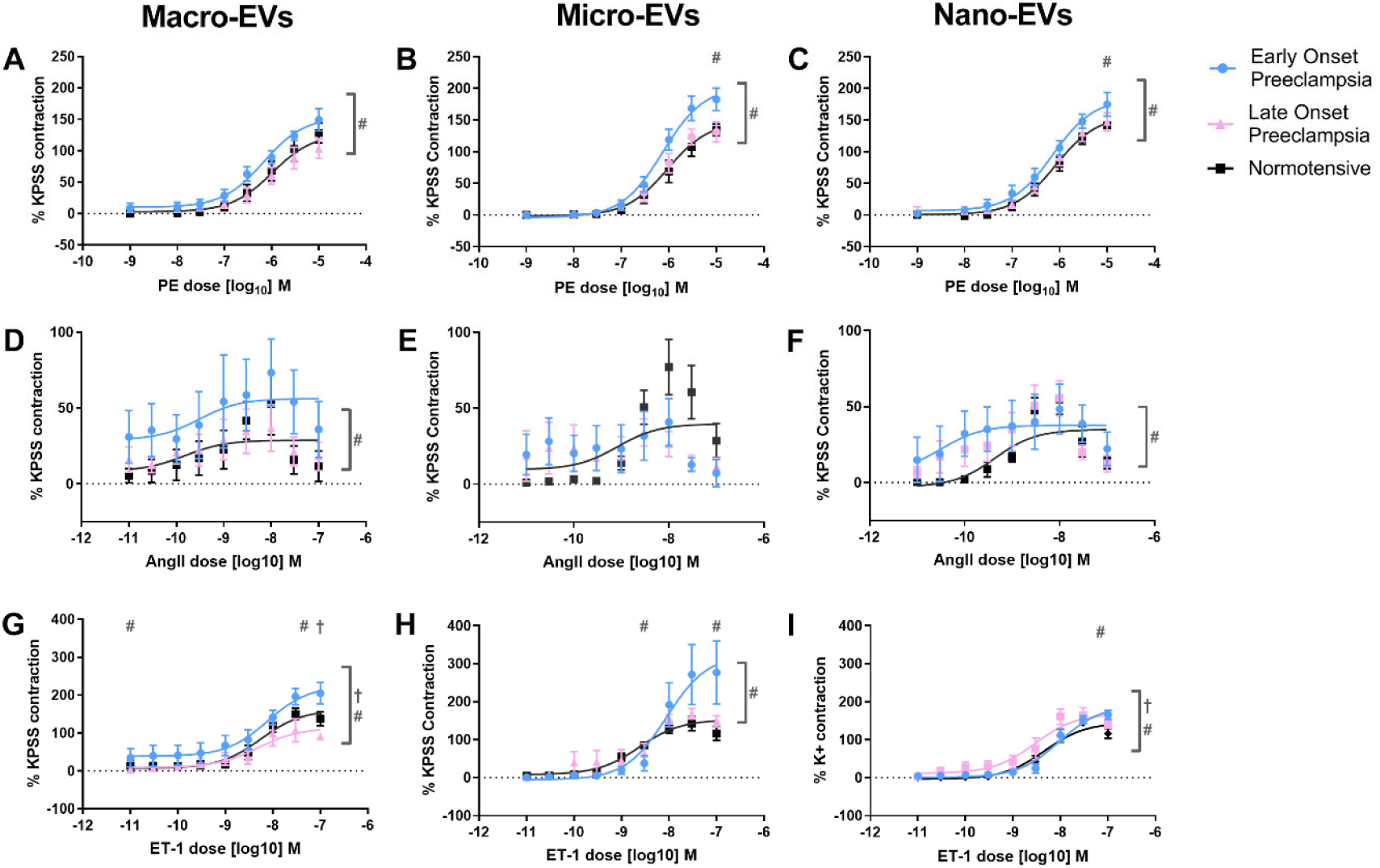
Responsiveness to the vasoconstrictors in mesenteric vessels from animals following 24 hours intravenous exposure to placental EVs from normal and preeclamptic pregnancies. Placental EVs from early-onset preeclampsia (blue circles) enhanced the constriction response to phenylephrine (PE) (A-C), angiotensin II (AngII) (D-F) and endothelin-1 (ET-1) (G-I) compared to EVs from normotensive pregnancies (black squares, # p<0.05). There was no difference in response to PE and AngII between late-onset preeclampsia (pink triangles) and normotensive pregnancy groups but nano-EVs from late-onset preeclampsia appeared to enhance the constriction in response to ET-1 († p<0.05) whilst macro-EVs appeared to enhance the constriction response to ET-1.

Placental macrovesicles and nanovesicles from pregnancies with early-onset preeclampsia also impaired both ACh and SNP mediated vasodilation (both p<0.05, blue circles, Figure 2) compared to vesicles isolated from normotensive pregnancies (black square, Figure 2).

**Figure 2.**
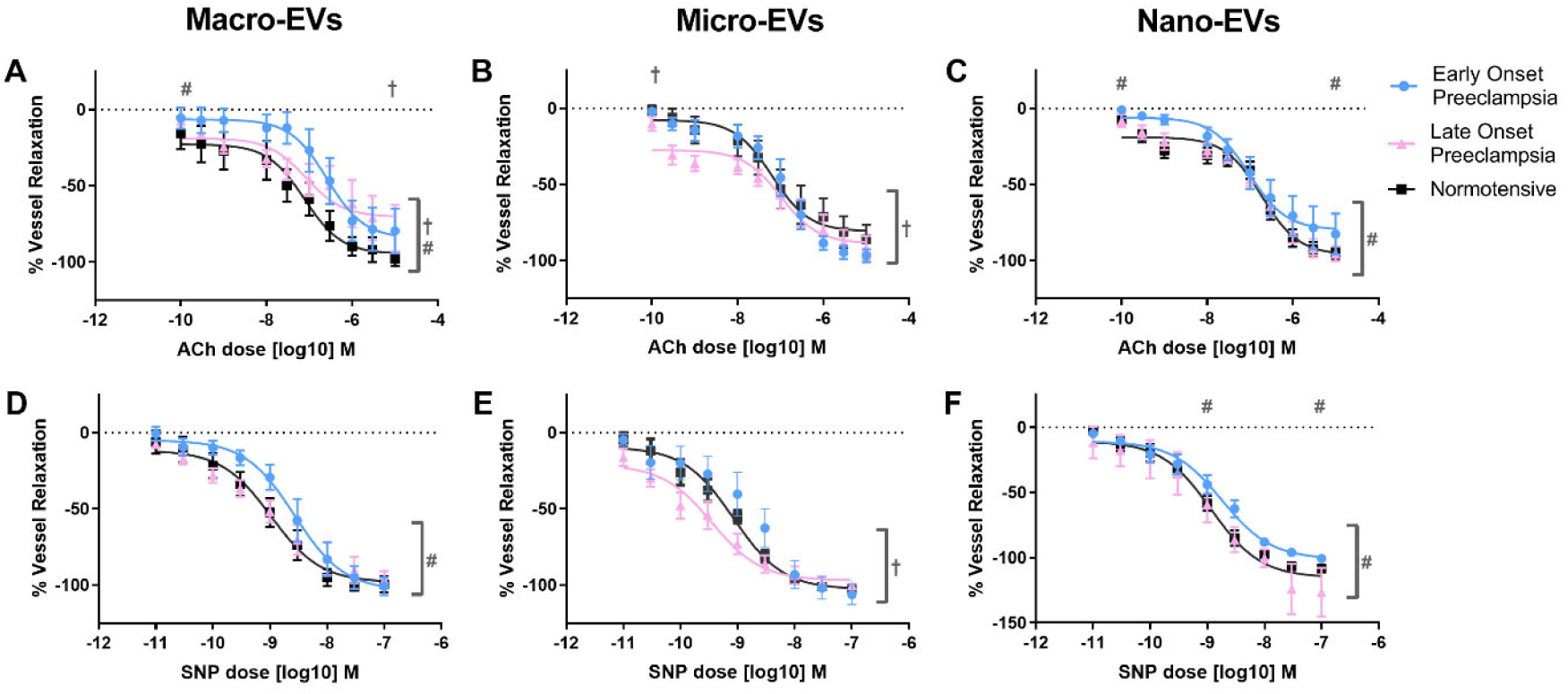
Responsiveness to vasodilators in mesenteric vessels from animals following 24 hours intravenous exposure to placental EVs from normal and preeclamptic pregnancies. Placental macro-EVs and nano-EVs from early-onset preeclampsia (blue circles) reduced the relaxation response to both acetylcholine (ACh) (A-C) and sodium nitroprusside (SNP) (D-F) compared to the normotensive group (black squares) (# p<0.05). Micro-EVs from late-onset preeclampsia (pink triangles) enhanced the vasodilation to both ACh and SNP (†p<0.05) whilst the macro-EVs appeared to reduce the vasodilation to ACh but not SNP.

Macrovesicles from pregnancies with early-onset preeclampsia impaired the Ach-mediated vasodilation response across the doses tested (p<0.0001) but almost abolished the vasodilation response to ACh at lower doses (p=0.0393).

#### Response to placental EVs from pregnancies with late-onset preeclampsia 24 hours after injection

Mesenteric arteries from mice injected intravenously with placental EVs from pregnancies with late-onset preeclampsia did not altered reactivity to the vasoconstrictors PE and AngII compared to mesenteric arteries from mice injected with placental EVs from normotensive pregnancies. The response to the vasoconstrictor ET-1 was mixed with macro-EVs from late-onset preeclampsia reducing the maximum generated response to ET-1 at higher doses, nano-EVs increasing the response to ET-1 across all concentrations, and micro-EVs not eliciting a different response (pink triangles, Figure 1) compared to EVs from normotensive pregnancies (black squares, Figure 1).

The response of mesenteric arteries from mice injected intravenously with placental EVs from pregnancies with late-onset preeclampsia to vasodilators was mixed. Micro-EVs enhanced vasodilation of mesenteric arteries to both ACh (p=0.0172) and SNP (p=0.0002), while macro-EVs restrained vasodilation in response to ACh (p=0.0057, pink triangles, Figure 2) but had no effect on vasodilation in response to SNP. Nano-EVs from pregnancies with late-onset preeclampsia did not alter the response to either ACh or SNP compared to normotensive pregnancies (black squares, Figure 2).

### Vascular responsiveness of resistance arteries 30 minutes after injection of placental EVs from pregnancies with preeclampsia or normotensive pregnancies

The contractile responses to AngII were increased in vessels that had been exposed to both placental micro-EVs (p=0.002) and nano-EVs (p=0.0228) from early-onset preeclampsia compared to normotensive pregnancies (Figure 3) 30 minutes after administration. However, in response to ET-1, both macro-EVs (p=0.0025) and nano-EVs from early-onset preeclampsia elicited a reduced contractile response to ET-1 compared to EVs from normotensive pregnancies. The response to PE was not statistically different after 30 minutes exposure to any type of placental EVs from early-onset preeclampsia compared to normotensive pregnancies.

**Figure 3.**
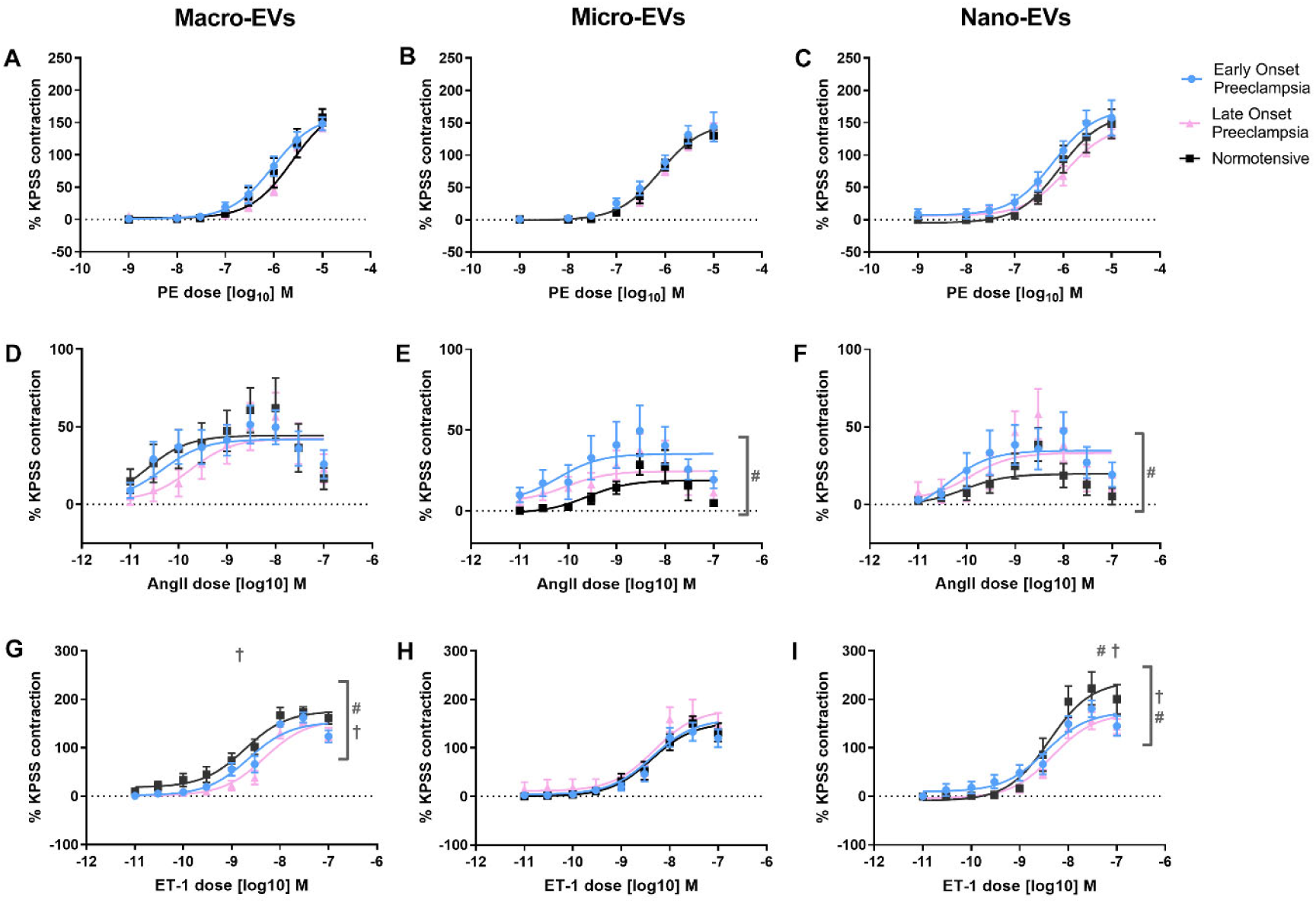
Responsiveness to vasoconstrictors in mesenteric vessels from animals following 30 minutes intravenous exposure to placental EVs from normal and preeclamptic pregnancies. There was no difference in the constriction response to phenylephrine (PE) (A-C) between EVs from early-(blue circles) or late-onset (pink triangles) preeclampsia and normotensive groups (black squares). Exposure to macro-EVs and nano-EVs from both early-onset (#, p<0.05) and late-onset (†, p<0.05) preeclampsia resulted in a reduced constriction to endothelin-1 (ET-1) (G-I). Only micro-EVs and nano-EVs from early-onset preeclampsia enhanced the constriction to angiotensin II (AngII) (D-F) compared to the normotensive groups (p<0.05).

All three placental EV types from pregnancies with early-onset preeclampsia reduced the relaxation of mesenteric resistance arteries in response to ACh (all p<0.05, Figure 4). Only placental nano-EVs from pregnancies with early-onset preeclampsia reduced the relaxation response to SNP compared to EVs from normotensive pregnancies 30 minutes after administration of the EVs (p=0.0004), almost abolishing the relaxation response at the lower doses (p<0.0001).

**Figure 4.**
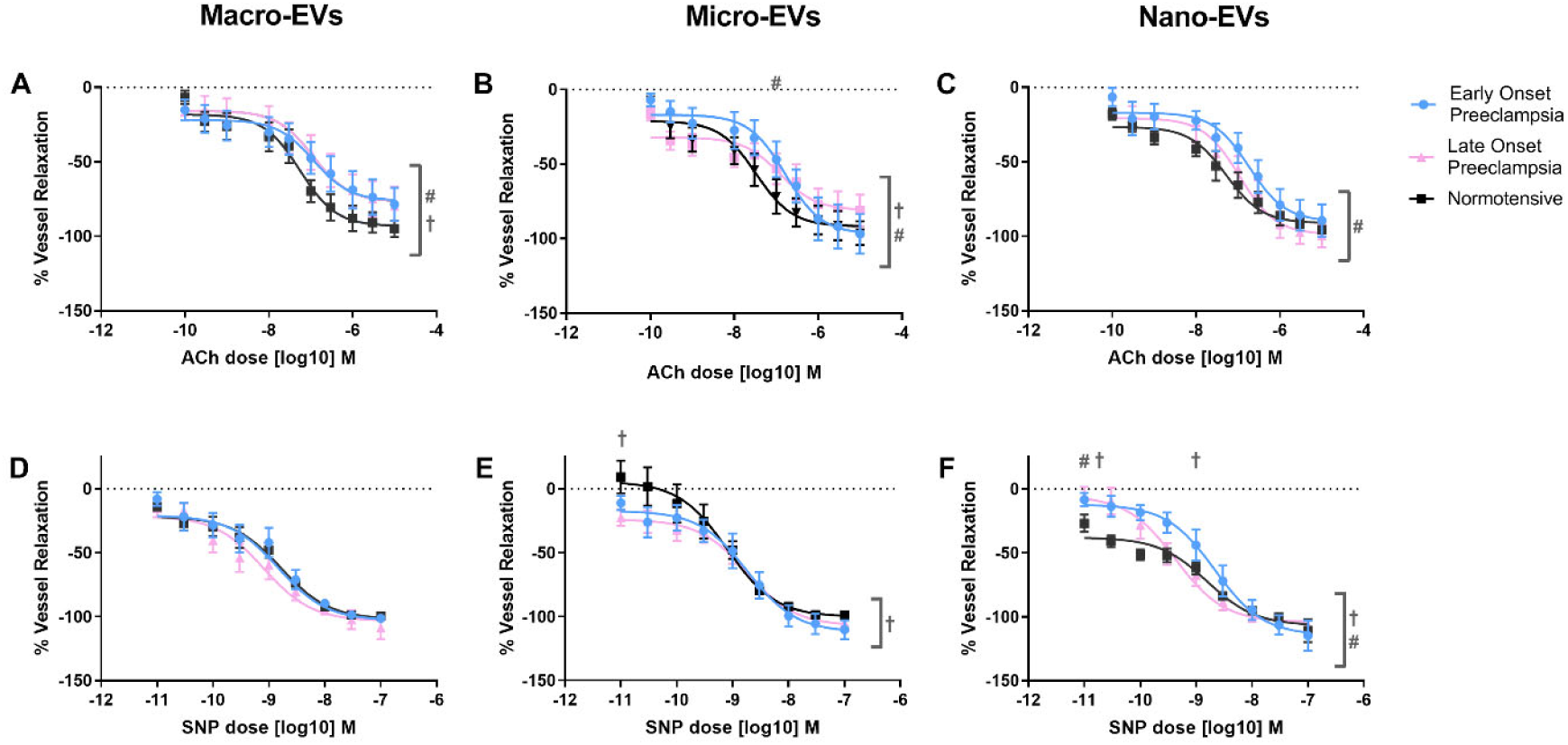
Responsiveness to vasodilators in mesenteric vessels from animals following 30 minutes intravenous exposure to placental EVs from normal and preeclamptic pregnancies. All three vesicles types from early-onset preeclampsia (blue circles) but only macro-EV and micro-EVs from late-onset preeclampsia (pink triangles) reduced the relaxation to acetylcholine (ACh) (A-C) compared to normotensive groups (black squares). Nano-Evs from early-onset preeclampsia reduced vasodilation in response to sodium nitroprusside (SNP) (#, p<0.05) (D-F) while in the late-onset preeclampsia groups, micro-EVs appeared to enhance the relaxation and nano-EVs reduce the relaxation response (†, p<0.05)

### Placental EVs from pregnancies affected by late-onset preeclampsia reduced mesenteric artery response to ET-1 compared to EVs from normotensive pregnancies 30 minutes after injection

In response to ET-1, both macro-EVs and nano-EVs unexpectedly reduced the contractile responses (p<0.0001 and p=0.0008, respectively, figure 3). Furthermore, macro-EVs reduced the sensitivity of the vessels to ET-1 with an altered EC50 (p=0.0225) and nano-EVs reduced the maximal contractile response at the higher doses (p=0.0063, figure 3). Placental micro-EVs from pregnancies with late-onset preeclampsia did not affect the reactivity to ET-1 compared micro-EVs from normotensive pregnancies. There was no effect on the vascular response to PE and AngII in vessels from mice 30 minutes post administration of placental macro-EVs, micro-EVs, or nano-EVs from pregnancies with late-onset preeclampsia compared to normotensive pregnancies.

Ach-mediated vasodilation was impaired in mesenteric arteries from mice injected with placental macro-EVs and micro-EVs, but not nano-EVs from pregnancies with late-onset preeclampsia compared to normotensive pregnancies (p=0.0075, p=0.0224 respectively, Figure 4). Whilst nano-EVs from pregnancies with late-onset preeclampsia restrained SNP-mediated vasodilation (p<0.0001), with an almost absent response to SNP at lower doses (p<0.0001) and an altered sensitivity (p=0.0069), micro-EVs appeared to enhance SNP mediated vasodilation (p=0.011) with a larger relaxation at lower doses (p=0.0001). Macro-EVs from pregnancies with late onset preeclampsia did not affect SNP mediated vasodilation.

### The effects of normotensive placental EVs on maternal vasculature were vesicle type specific and changed between 30 minute and 24 hours

To determine whether the time-dependent differences observed in the effects induced by early-onset preeclampsia were due to an alteration in the response to the preeclampsia-derived placental EVs, or due to the changes in our comparison group – placental EVs from normotensive pregnancies, we compared the 30 minute or 24 hour groups injected with placental EVs from normotensive pregnancies.

The response to AngII were vesicle type-specific, with micro-EVs (p<0.0001) and nano-EVs (p=0.0260) enhancing the response to AngII at 24 hours compared to 30 minutes, but macro-EVs reducing the response to AngII (p=0.0473) (Figure 5). Both macro-EVs (p<0.0001) and nano-EVs (p<0.0001) reduced the response to ET-1 at 24 hours, but micro-EVs did not exhibit a time-dependent response (Figure 5). No differences in the response to PE were observed between the 30 minute and 24 hour circulation times.

**Figure 5.**
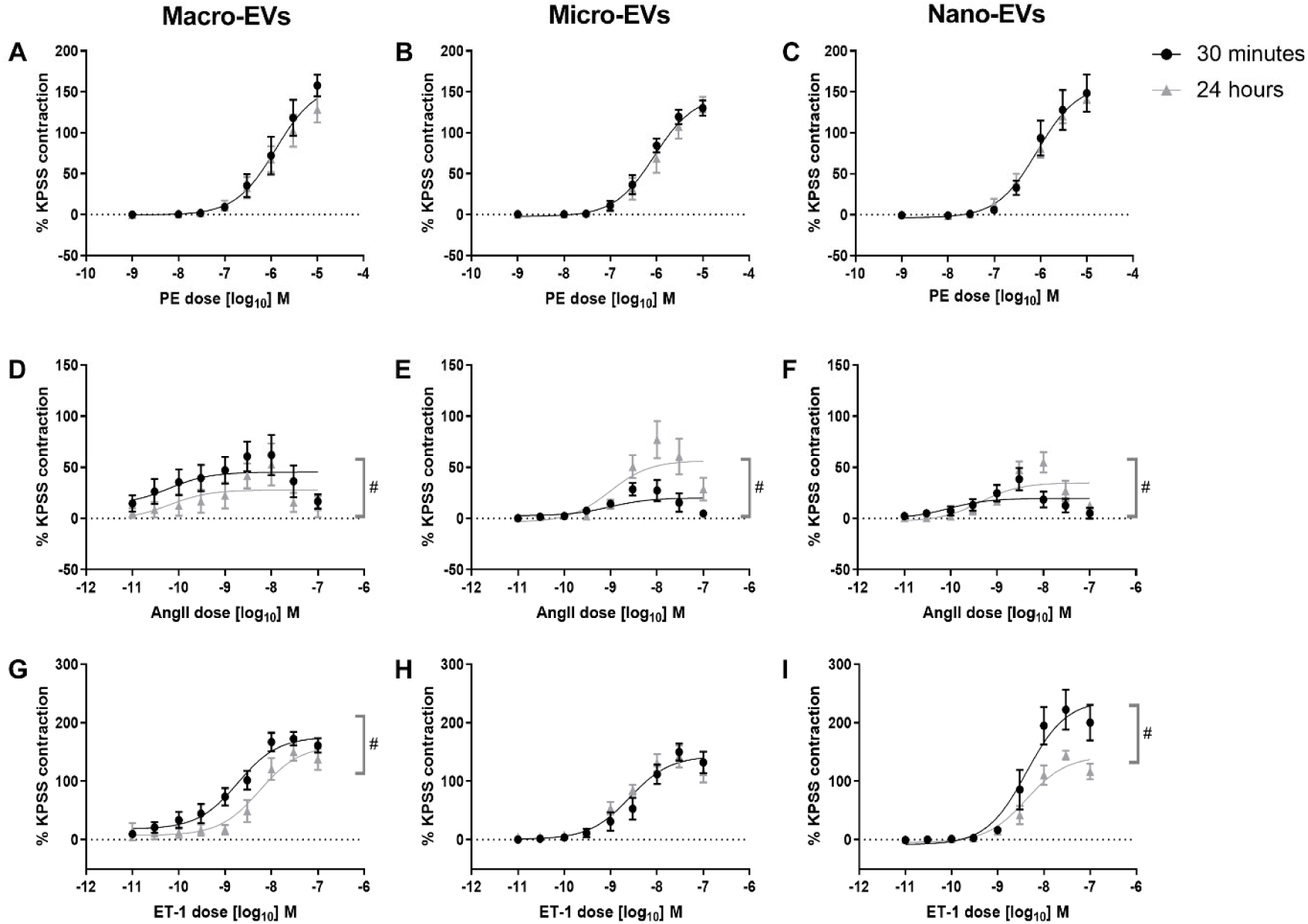
Time dependent effects to vasoconstrictors in vessels from mice exposed to placental EVs from normotensive third trimester pregnancies. The contraction to angiotensin II (AngII) (D-F) was enhanced at 24 hours (grey triangles) in the micro-EV and nano-EV groups compared to the corresponding response at 30 minutes (black circles) (#, p<0.05). In contrast, in the macro-EVs groups, the response was reduced. Similarly the response to endothelin-1 (ET-1) (G-I) was reduced at 24 hours in the macro-EV and nano-EV groups (p<0.05).

Micro-EVs (p=0.002) and nano-EVs (p<0.0001) restrained vasodilation in response to ACh at 24 hours compared to 30 minutes while nano-EVs restrained vasodilation in response to SNP (p<0.0001) at 24 hours (Figure 6).

**Figure 6.**
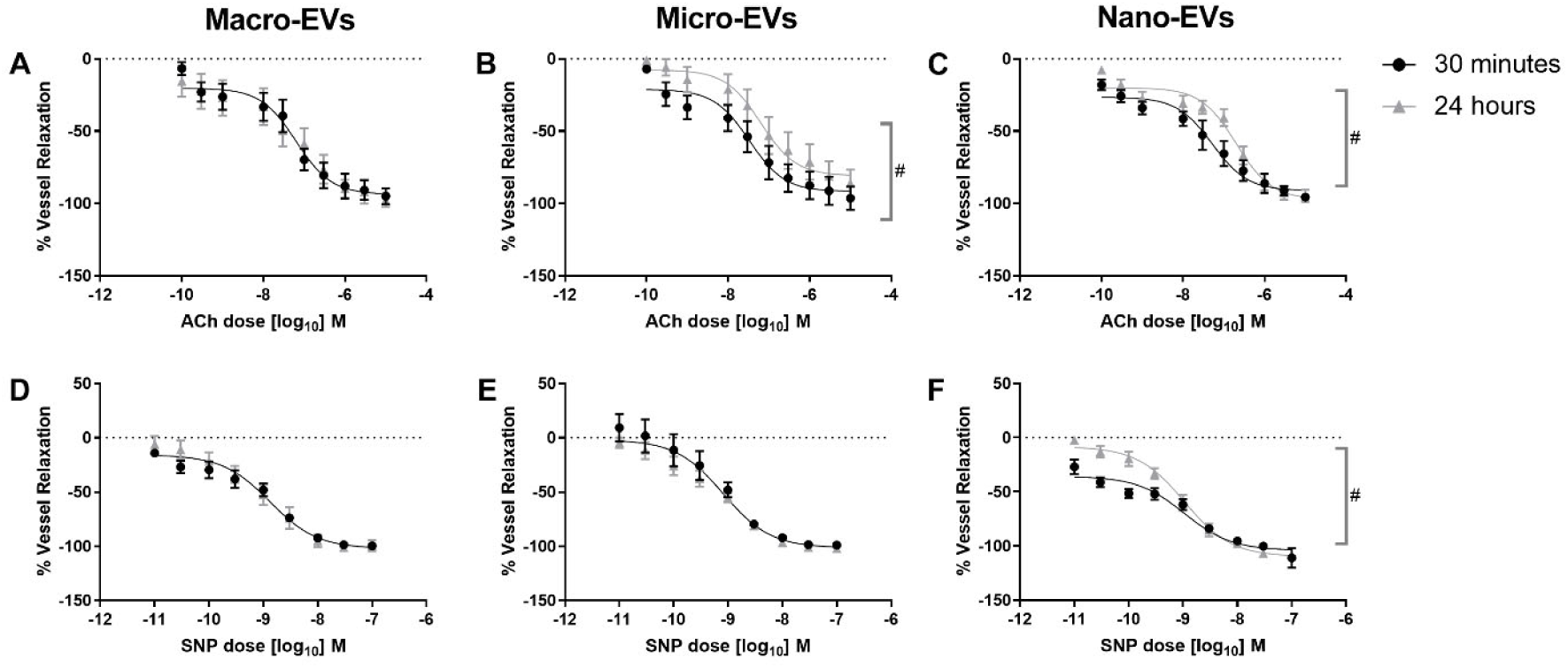
Time dependent effects of vasodilators in vessels from mice exposed to placental EVs from normotensive third trimester pregnancies. Exposure to micro-EVs for 24 hours (grey triangles) reduced the response to acetylcholine (grey triangles) (A-C) compared to 30 minutes exposure (black circles) (#, p<0.05). Similarly, exposure to nano-EVs for 24 hours reduced the relaxation response to sodium nitroprusside (SNP) (D-f) (#, p<0.05).

## Discussion

We confirmed our hypothesis that placental EVs have the potential to play a direct role in modulating the increase in vascular resistance in women with early-onset preeclampsia. We demonstrated that the placental EVs from early-onset preeclampsia enhanced vasoconstriction and reduced vasodilation in mesenteric arteries 24 hours after injection. In contrast, this pro-constrictive phenotype was not observed in mesenteric arteries from mice injected with placental EVs from late-onset preeclampsia. We also demonstrated that the placental EVs from normotensive term human pregnancies injected i.v. into pregnant mice had a time-dependent component to the response of mesenteric arteries using wire myography *ex vivo*, similar to our previous results after injecting first trimester EVs (10, 34). However, the changes in the responses to normotensive EVs did not explain the time dependent changes in the response to preeclamptic EVs.

We have previously published that human placental EVs from first trimester placenta can modulate the response of small resistance arteries in pregnant mice (34), potentially contributing to the decrease in vascular resistance that is an essential central component of the normal maternal cardiovascular adaptation to pregnancy (35). This study demonstrates that human placental EVs from third trimester pregnancies also have the potential to alter the responsiveness of resistance arteries to vasoactive stimuli.

A hallmark of preeclampsia is the alteration of endothelial cell function towards a pro-thrombotic, pro-inflammatory and pro-constrictive state (endothelial dysfunction) (36). In this study, we report that placental EVs from pregnancies with early-onset preeclampsia can also induce a pro-constrictive state in small resistance arteries studied *ex vivo*, from as soon as 30 minutes after injection. Twenty-four hours after injection, placental EVs from early-onset preeclampsia altered the function of small resistance arteries (both endothelial cells and surrounding smooth muscle) towards a pro-constrictive and anti-vasodilatory state. In contrast, placental Evs from pregnancies with late-onset preeclampsia did not have this same effect. These findings suggest firstly that the content of the placental EVs is functionally different between early– and late-onset variants of preeclampsia (as the dose of EVs was the same). Secondly, they suggest that placental EVs alone may be sufficient to induce a pro-constrictive, anti-vasodilatory vascular reactivity in women with early-onset preeclampsia. Alternatively placental EVs may contribute to this altered vascular tone alongside other vasoactive substances thought to be important in the pathogenesis of preeclampsia. These may include altered nitric oxide or endothelin-1 pathways which can act downstream to factors released from placental ischemia or those altering angiogenic balance [e.g. sFlt-1, reviewed in (37)). Finally, placental EVs from late-onset preeclampsia do not produce, or alone are insufficient to alter vascular reactivity towards a pro-constrictive phenotype.

### The effect of placental EVs mirrors maternal haemodynamic profiles from early-onset and late-onset preeclampsia

Maternal hemodynamic profiles differ between women affected by early-onset preeclampsia, late-onset preeclampsia, and normotensive pregnancies (reviewed in 21). Women with early-onset preeclampsia have higher systemic vascular resistance throughout pregnancy and a lowered cardiac output (38, 39). In contrast, women with late-onset preeclampsia have lower systemic vascular resistance and elevated cardiac output (38). The ability of placental EVs from early-onset preeclampsia to enhance arterial constriction in response to vasoconstrictors and impair reactivity to vasodilators, while placental EVs from late-onset preeclampsia did not, aligns with the haemodynamic profiles of the two presentations of preeclampsia suggesting that placental EVs may be a key driver of these vascular changes in preeclampsia, particularly early-onset preeclampsia. Furthermore, these data further support the current view that the underlying aetiology and pathogenesis is different between early– and late-onset preeclampsia, and that early-onset preeclampsia is driven primarily by a pathologic placenta (40, 41). In contrast, late-onset preeclampsia results primarily from maternal maladaption to the physiologic needs of pregnancy (40, 41).

### Comparison of effects to the exposure to plasma EVs vs isolated placental EVs from preeclamptic pregnancies

To the best of our knowledge, this is the first comparison of vascular function after exposure to the full range of EVs produced by placenta from early-onset and late-onset preeclampsia and normotensive human placentae. Previous studies comparing EVs from normotensive pregnancies and preeclampsia focused on studying the vascular responses to maternal plasma EVs (42–45) but placental EV comprise only 10-15% of plasma EVs in pregnant women (46). A major difference exists between our work and existing literature as maternal plasma contains a large mixture of EVs, including those from platelets (47), endothelial cells (48, 49), lymphocytes (50) and from other organs (51) and the EVs from these cell types are known to impair vasodilation independent of placental EVs (48–52). Furthermore, the cocktail of EVs isolated from maternal plasma represents a mixture of both the triggering placental EVs, as well as the EVs resulting from the maternal pathological (or physiological) response to the placental EVs and those EVs that from the maternal response may have effects that either enhance or inhibit the effects of placental EVs. For example, a previous study treating resistance arteries of normotensive Wistar Kyoto rats with plasma-derived EVs from spontaneous hypertensive rats demonstrated that these plasma EVs can ‘encode’ a compensatory shift in the response to ACh-mediated vasodilation after the development of hypertension in the animals compared to EVs taken from plasma before hypertension develops (53). These data combined suggest that to understand the role of placental EVs in triggering preeclampsia, it is essential to avoid the confounding effects of non-placental EVs as we have done in this work. Additionally, other studies have used experimentally damaged human or murine placentae that may not reflect the EVs produced by placentae from pregnancies with preeclampsia (54, 55) or may have focused on a single EV type (54).

### The response of maternal vasculature to placental EVs is time dependent

This study demonstrates that the effect of placental EVs from normal pregnancies and pregnancies with preeclampsia was different at 30 minutes and 24 hours after injection. Time-dependent effects of placental EVs have been demonstrated previously by both our lab (10) and others (45). This time-dependent effect on maternal vascular reactivity appears to be a feature of placental EVs as circulating maternal cell-derived EVs do not have this time-dependency. For example, plasma EVs, endothelial cell-derived EVs or blood-cell derived EVs have been demonstrated to impair endothelium-dependent vasodilation (2–9) but endothelial cell-derived EVs impair acetylcholine mediated vasodilation regardless of whether the effect was measured 30 minute (5), 3 (4) or 24 hours after exposure (56).

Whilst this study uncovered some interesting time-dependent functional differences of placental EVs from pregnancies with early-onset and late-onset preeclampsia, it also leads to questions to how these functional differences arise. Studies have previously compared the proteome of placental EVs from pregnancies with preeclampsia with normotensive pregnancies and found unique proteins or increase protein abundance in placental EVs from pregnancies with preeclampsia (25, 26). For example, preeclamptic EVs carry more Flt-1 than normotensive placental EVs (57, 58) and it may be that the 30 minutes response is dominated by proteins carried by the preeclamptic EVs. However, the altered response at 24 hours compared to 30 minutes suggests that the EVs may have delivered signals required new gene expression in maternal cells, or that in addition to proteins, other bioactive molecules may be involved in the development of these effects, such as mRNA or miRNA. It has been published that miRNA is present in placental EVs, which can be transferred into endothelial cells with functional activity *in vitro* (17). Furthermore, women affected by preeclampsia have higher expression levels of a number of miRNAs in the placenta compared to women with normotensive pregnancies (59). It could be possible that the shift in our observed effects at 24 hours could be in part induced by miRNAs, although further study is required to establish this. We stress the importance for future studies to evaluate functional effects of placental EVs consistently at a single time point, or preferably that new studies evaluate a range of time points.

### The effect of placental EVs on maternal resistance arteries is specific to the EV type

One of the strengths of this work is that we investigated the effect of placental macro-, micro– and nano-EVs individually to determine the effect of each type of vesicle on vascular reactivity. We have previously reported that different EV types from first trimester placenta contain different protein cargoes (60) and therefore may affect the maternal vasculature differently. It is likely that these differences also exist in the third trimester EVs, and is reflected in the functional differences between the three types of EVs studied here, regardless of whether the EVs were from pregnancies that were normotensive or with preeclampsia. However, this also creates limitations in our interpretation of the data as the human placenta simultaneously releases a combination of the three different types of placental EVs. Given the differential effects of each vesicle type, we do not yet know when delivered in combination, as occurs in pregnancy, what the overall effect of the EVs would be. It is also unclear whether the proportion of each vesicle type released is altered in preeclampsia. The differential effects of vesicle types on maternal vasculature also potentially poses a question on whether researchers should be focused on the total number of EVs released from the placenta in pathological pregnancies, or is it the differential proportion of each released vesicle type that leads to maternal pathology? This is especially true since the amount of EVs released from the placenta is drastically increased in preeclampsia. Here, we have clearly shown by delivering equal amounts of EVs that these different responses to these EV types are due to differences in the content of the EVs rather than different amounts of the EVs.

### Can we translate this work to human pathology?

One limitation of this work is the use of a cross-species model in which human placental EVs were injected into pregnant mice. However, preeclampsia is essentially unique to humans; rodents do not exhibit preeclampsia nor do they not produce macro-EVs which have been associated with the pathogenesis of preeclampsia (61). Although the use of placental EVs from mice would overcome any potentially xenogenic effects, we cannot truly study the effects of preeclampsia on the placenta using mouse placentae. Similarly, it would have been possible to study the effect of human EVs on isolated human omental or other arteries. However, such a study would be entirely *ex vivo* and could not account for features that require a whole animal such as the natural biodistribution of EVs, and such studies cannot look at longer time points.

Whilst we have observed alterations in vascular reactivity, this does not translate directly to altered systemic vascular resistance as there are other mechanisms which control vascular resistance such as flow and sheer stress responses, autonomic control as well as various other vasoactive substances not investigated in this work. Our results point to a crucial role of placental EVs in the development of preeclampsia but further investigations using more complex integrated physiological models will be required to confirm the role of placental EVs in the pathogenesis of preeclampsia.

### Perspectives

The development of the hypertensive disease of pregnancy, preeclampsia, involves a trigger from the placenta which induces endothelial dysfunction – a hallmark of the disorder. Here, we report that extracellular vesicles (EVs) isolated from the placenta of pregnancies with early-onset preeclampsia, but not late-onset preeclampsia, induce a pro-constrictive, anti-vasodilatory effect on resistance arteries after 24 hours circulation *in vivo* in the pregnant mouse. This suggests that in early-onset preeclampsia, placental EVs may be the factor released from the placenta that triggers the development of maternal disease. However, the response of the vasculature to vasoconstrictive and vasodilatory substances is only one aspect of the alterations in the maternal circulatory system resulting in preeclampsia. In future studies, it would be worthwhile to investigate the full hemodynamic profile of rodents injected with preeclamptic placental EVs to determine whether placental EVs alone are enough to induce the plethora of hemodynamic changes seen in pregnancies complicated by preeclampsia.

### Novelty and Relevance

*Novelty*: These data demonstrate for the first time that isolated human placental EVs from women with early-onset preeclampsia were able to produce a pro-constrictive, anti-vasodilatory state in the resistance arteries of pregnant mice, compared to placental EVs from normotensive pregnancies. It was also demonstrated that this effect was induced only by early-onset, and not by late-onset preeclamptic placental EVs. *Relevance*: Eliciting these altered responses in resistance arteries may directly contribute towards the development of the high-resistance state and hypertension in women with preeclampsia.

## Acknowledgments

The Authors wish to thank the women and whānau who generously donated their placenta for this study, the Labour and Birthing Unit staff at Te Whatu Ora Te Toka Tumai Auckland, and Ms Rebecca Hay, Ms Mariska Oakes-Ter Bals and Ms Laura Mackay for their support in the recruitment of participants and collection of placenta and clinical data.

