## Supplementary Figures and Methods for "Placental extracellular vesicles from early-onset but not late-onset preeclampsia induce a pro-vasoconstrictive and anti-vasodilatory state in resistance arteries"

**Supplementary Table 1. Maternal and infant characteristics of the pregnancies included in this study. As expected, systolic and diastolic arterial pressures were higher in the preeclampsia groups compared to the normotensive group.** Gestational age at delivery, infant birthweight and placental weights at delivery were also lower in the early-onset preeclampsia group compared to the late-onset preeclampsia group and the normotensive group. Maternal age, body mass index (BMI) and number of nulliparous women included were comparable between groups. # denotes significant difference ($p<0.05$) between early-onset and late-onset preeclampsia, † denotes significant difference ($p<0.05$) of early-onset or late-onset preeclampsia compared to normotensive groups.

| | **Early-onset preeclampsia (n=8)** | **Late-onset preeclampsia (n=9)** | **Normotensive pregnancy (n=13)** | ***p-valued compared to late-onset preeclampsia** | **†p-valued compared to normotensive** |
|---|---|---|---|---|---|
| **Maternal characteristics ((median [IQ range])** | | | | | |
| Age | 33.5 [30.25 - 37.75] | 34.0 [27.50 - 36.00] | 35.0 [31.00 - 36.50] | ns | ns |
| BMI | 25.30 [22.03 - 34.28] | 24.20 [20.50 - 30.50] | 20.95 [20.00 - 27.30] | ns | ns |
| Nulliparous | 5/8 | 8/9 | 4/13 | n/a | n/a |
| Systolic blood pressure (mmHg) | 165 [156 - 172]† | 146 [138 - 163]† | 118 [108 - 124] | ns | **p<0.01** |
| Diastolic blood pressure (mmHg) | 102 [94 - 108]† | 97 [91 - 102]† | 75 [72 - 81] | ns | **p<0.001** |
| Antihypertensive medicine prescribed | 6/8 | 4/9 | n/a | n/a | n/a |
| **Infant characteristics** | | | | | |
| Gestational age at delivery (days) | 227 [217 - 261]*† | 264 [261 - 276] | 274 [273 - 279] | **0.0164** | **0.0033** |
| Infant birthweight (grams) | 1500 [1120 - 3440]*† | 2780 [2630 - 3345] | 3520 [3300 - 3800] | **0.0278** | **0.0002** |
| Placental weight (grams) | 305 [225 - 495]*† | 495 [465 - 580]† | 655 [548 - 678] | **0.0286** | **0.0261** |
| Customised birthweight centile ≤1% | 5/8 | 2/9 | 0/13 | n/a | n/a |
| Male infant | 3/8 | 5/9 | 8/13 | n/a | n/a |
| Number of live births | 8/8 | 9/9 | 13/13 | n/a | n/a |

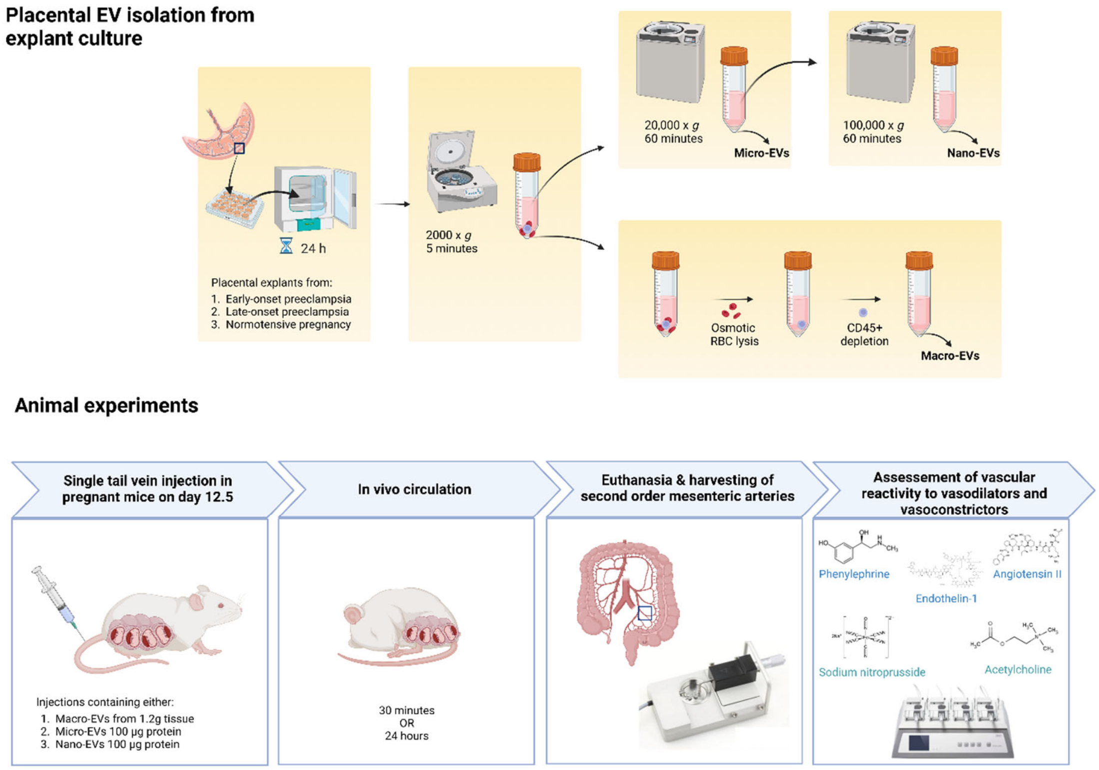

**Supplementary Figure 1. Workflow of placental explant culture model used to isolate placental EVs and animal experiments.** Explants of placental villous tissue from early-onset preeclampsia, late-onset preeclampsia, and normotensive pregnancies were incubated for up to 24 hours to collect the explant medium. The conditioned medium went through sequential centrifugation steps (2,000 x *g*, 20,000 x *g* and 100,000 x *g*) to collect the macro-EVs, micro-EVs and nano-EVs respectively. The macro-EV pellet underwent osmotic lysis to remove contaminating red blood cells and CD45 depletion using anti-CD45 magnetic beads to remove any CD45+ cells. Each EV type from early-onset preeclampsia, late-onset preeclampsia and normotensive pregnancies were injected separately into individual mice and allowed to circulate for either 30 minutes of 24 hours before sacrificing and removal of second order mesenteric arteries for assessing reactivity to vasoactive substances. There were 18 experimental groups in total.

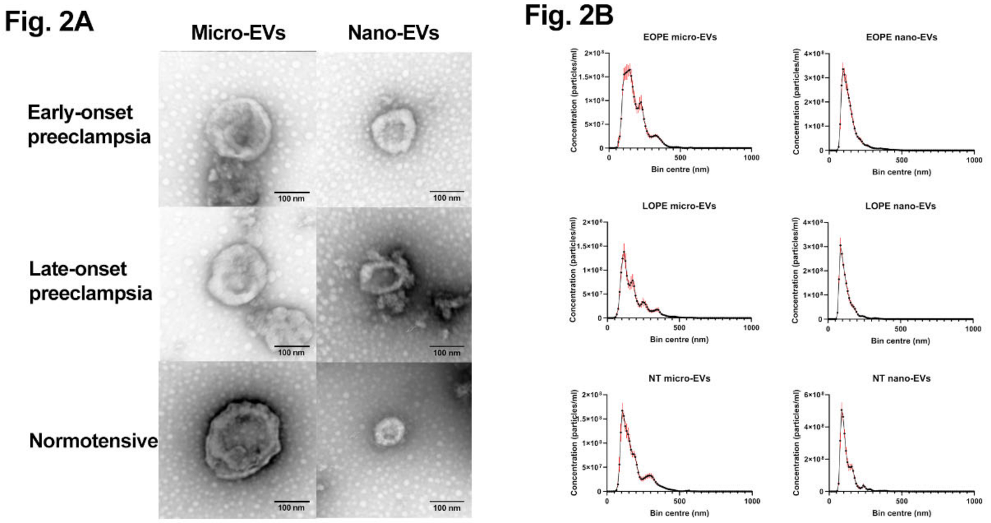

**Supplementary Figure 2.** Transmission electron micrographs (1A) and nanoparticle tracking analysis (1B) of placental microvesicles (micro-EVs) and nanovesicles (nano-EVs) from early-onset preeclampsia (EOPE), late-onset preeclampsia (LOPE) and normotensive pregnancies (NT). The modal sizes and appearances under transmission electron microscopy were not different between the groups.

**Supplementary Methods. Nanoparticle tracking analysis and transmission electron microscopy imaging of extracellular vesicles**

For nanoparticle tracking analysis (NTA), EVs were diluted at 1:100 in PBS and three 30 second videos were taken using a Nanosight NS300 (Malvern Panalytical) under low flow conditions (Screen gain:1, Camera level:14) and characterized using the Nanosight 3.4 software (Screen gain:10, Detection threshold:6). For transmission electron microscopy (TEM), EV samples were transferred from PBS buffer to ultrapure water by loading roughly 200 µl of the EVs into Vivaspin 500 (Sartorius AG) centrifugal concentrators with a 100 kDa cutoff and centrifuging at 10,000 x*g* until most of the PBS had flowed through the filter (roughly 10 minutes).  450 µl of ultrapure water was then added to the filter and the centrifugation process was repeated twice more before finally resuspending the EVs in 100 µl ultrapure water. Negative staining was conducted by first adsorbing EVs onto Formvar-coated copper grids (Electron Microscopy Sciences) for 10 minutes. Excess liquid was carefully removed with filter paper (Whatman) and the copper grid was then transferred to 20 µl filtered uranyl acetate for 2 minutes. Excess liquid was again removed with filter paper and the grid was allowed to dry under a lamp for 10 minutes. Grids were visualised on a Tecnai G2 Spirit TWIN (FEI, Hillsboro, OR, USA) TEM at 120 kV accelerating voltage, and images were captured using a Morada digital camera (SIS GmbH, Munster, Germany).
